# Discovering Novel Therapeutic V_H_Hs for Emerging Viruses: Perspectives from VEEV Selection Strategies

**DOI:** 10.64898/2026.01.24.701463

**Authors:** Autumn T. LaPointe, Fortunato Ferrara, Jennifer M. Zupancic, Alba L. Montoya, Jurgen Schmidt, Li-Wei Hung, Alison M. Kell, Nileena Velappan

## Abstract

Evolution or emergence of a new viral variant is a significant public health concern. Alphaviruses, such as Venezuelan equine encephalitis virus (VEEV), are mosquito-borne viruses which are becoming more prevalent due to expansion of vector habitats. The increased prevalence of such viruses provides opportunities for novel variants to evolve. Key therapeutic molecules that could be developed against viral pathogens are recombinant antibodies or antibody fragments, such as the variable heavy domain of heavy chain antibodies (V_H_Hs). These proteins can neutralize or sequester viral particles, preventing or reducing infection. However, due to the evolution of viruses, there is a need to isolate new antibodies and direct their binding to particular epitopes on the virus. *In vitro* selections offer a promising pathway for the selection of therapeutic antibodies, but as we demonstrate, the choice of a target for these selections is key to obtaining the desired viral binding characteristics. Here we report four novel “human” V_H_Hs which bind to the VEEV E2 protein selected using different strategies that include both computational and biochemical design of suitable antigens and whole virus selections. These V_H_Hs have distinct complementarity-determining regions (CDRs). Multiple V_H_Hs bind to the VEEV viral particles in ELISAs, and we report the peptide epitope recognized by these V_H_Hs. Though non-neutralizing, when immobilized, these V_H_Hs bind to and sequester VEEV viral particles preventing infection, demonstrating the potential of these V_H_Hs to perform viral “sponging”. The selection strategies we report may have applications to further antibody developments against other viruses.

**Significance/Importance:** Alphaviruses, and in particular Venezuelan equine encephalitis virus (VEEV), are recognized for their ability to cause severe disease and for their potential to be used as a biothreat. Despite this, there are currently no antiviral therapies or FDA-approved vaccines available to treat or prevent VEEV infection. This study reports on a novel antibody selection pipeline to produce antibody fragments against VEEV. Antibodies produced via this method showed strong affinity and high specificity to the VEEV E2 glycoprotein in multiple conformations. Additionally, while not neutralizing, the antibody fragments described were shown to be effective as “viral sponges”, having the ability to bind, sequester, and remove VEEV virions from solution, which represents a novel therapeutic approach.

## Introduction

Emerging pathogens with pandemic potential pose a serious threat to global public health. These viruses may emerge through natural evolution and zoonosis transmission or may be potentially released into susceptible populations either accidentally (e.g., laboratory leaks) or intentionally. Mosquito-borne viruses represent a particularly concerning risk in these scenarios due to socioeconomic conditions in tropical regions and the ongoing expansion of mosquito habitats driven by climate variations. Consequently, the development of novel therapeutics and innovative methods for generating these molecules is critical to safeguarding public health.

Alphaviruses are positive-sense, single-stranded RNA arboviruses capable of causing severe disease. Their natural enzootic transmission cycle involves a mosquito vector and mammalian or avian hosts; however, epizootic spillover events can lead to infection in humans and equids. Alphaviruses are broadly classified based on disease outcome as either arthritogenic or encephalitic. Arthritogenic alphaviruses, such as Chikungunya virus (CHIKV) and Ross River virus (RRV), cause illnesses ranging from mild febrile episodes to severe, persistent polyarthralgia that can last weeks to years’ post-infection.^1,2,3^ In contrast, encephalitic alphaviruses, such as Venezuelan equine encephalitis virus (VEEV), produce wide range of neurological symptoms, including potentially fatal encephalitis.^3,4^ VEEV is of particular concern as biodefense threat due to its high infectivity and potential for aerosol transmission.^5,6^ Despite the significant threat to public health, there are currently no FDA-approved vaccines or antiviral therapies available for prevention or treatment.

Antibody-based immunotherapy has proven to be a powerful tool against infectious diseases as demonstrated recently during the Coronavirus disease 2019 (COVID-19) pandemic.^7-9^Antibodies are highly effective therapeutics due to their target specificity, structural adaptability, and diversity.^9^ The variable heavy domain of heavy chain antibodies (V_H_Hs), also referred to as nanobodies, retain the binding ability of an antibody with a single protein domain and offer several advantages over conventional IgGs, including enhanced solubility, simplified engineering^10^, and superior stability across a broad pH range.^11-13^

Moreover, V_H_H domains are easily engineered in combination with other antibody formats. For instance, IgGs fused to additional V_H_H domain(s) have been actively pursued as therapeutics because they enable binding to multiple epitopes.^14^ Recently, Mei et al.^10^ described V_H_Hs developed against various viruses as well as currently available V_H_H fusion proteins. In their recent review on V_H_H antibody discovery and potential application, Alexander et al.^11^ denotes several potential therapeutic applications for “human” or humanized V_H_H antibodies in cancer biology, neurodegenerative diseases, autoimmune diseases, and infectious diseases. These publications highlight the unique niche for V_H_Hs as protein domains that can be tethered to other molecules to facilitate multi-specificity and the corresponding multifunctionality.

*In vitro* selections have been widely applied to the selection of anti-viral V_H_Hs due to the control provided over both the target antigen and the composition of V_H_Hs within the immune or synthetic library interrogated for binding.^15^ Synthetic V_H_H libraries that use human variable heavy chain as the scaffold for inserting Complementarity-Determining Regions (CDRs) allow for construction of V_H_H libraries.^16^ This design aims to address “humanness”, stability, and manufacturing issues.^17^ Libraries based on therapeutic V_H_H scaffolds that have been extensively studied and used in clinical trials provide frameworks that are well behaved with excellent developability properties.^18^ If the library and consequently the selected V_H_Hs contain human CDRs, this further facilitates their potential use as therapeutics.

V_H_H-based therapeutics have not been reported for VEEV. More importantly methods to select therapeutic V_H_H against emerging viruses that have complex viral coat proteins remain challenging. In the past couple of decades, *in vitro* binder (antibodies, peptides, other engineered molecules) selections using phage and yeast display have resulted in high quality moieties that recognize their intended target with high specificity and affinity.^9,19-22^ However, isolation of high quality binding proteins depends on the presentation of the antigen during these selections. Therefore, antigen formats that can be used in binder selection and/or evolution are key for developing novel therapeutics against emerging viruses.

Transmembrane coat protein ectodomains in viral particles are often solvent accessible or immunogenic and hence present the best antigen for *in vitro* or *in silico* binder selection and verification. However, recombinant expression and purification of these ectodomains for binder selection may be challenging for some viruses, especially if the coat protein is a heteromultimer. The ectodomain of a given protein may not be expressible or foldable in bacterial, mammalian, or cell free systems. Use of virus-like particle (VLP, pseudovirus) based protocols require extensive optimization and remain expensive. An alternative option is to use linear peptides from the viral coat protein as a selection antigen, followed by verification on infected cells or virus particles. *In vitro* antibody fragment selection on inactivated virus is also possible.

VEEV presents a unique opportunity to explore various antigen formats to select therapeutic V_H_H antibodies against a mosquito-borne pathogen with high pandemic potential. The major coat protein of VEEV is the heteromeric complex of three envelop glycoproteins E1/E2/E3.^23^ The availability of various fragments of this virus allowed us to explore *in vitro* V_H_H antibody selection using multiple antigen formats: 1) recombinant E2, 2) linear peptides of E2, selected based on molecular dynamics analysis, and 3) UV inactivated virus. Here we report on both the potentially therapeutic V_H_Hs discovered using these targets as well as the strategies and challenges associated with each of these antigen formats.

## Results

### Antigen selection

The availability of both recombinantly produced viral proteins and a crystal structure of the VEEV^23^ coat proteins enabled us to explore three antigen formats in our V_H_H selections (**Fig. 1a):** (1) recombinant proteins, (2) viral peptides, and (3) inactivated viruses. The purified recombinant ectodomain of E2, purchased from Creative Diagnostics, was initially chosen as the primary antigen. This small protein domain represents the dominant epitope targeted by previously reported antibodies.^24^ The antigen was dissolved in 1× PBS containing 8 M urea, as per the manufacturer’s instructions, and subsequently dialyzed into PBS prior to biotinylation. The requirement for urea raised concerns about protein stability in PBS during phage and yeast sorting. Structural analysis revealed the intertwined nature of the E1/E2 complex, indicating challenges could arise if regions of E2 that are not solvent exposed in the E1/E2 complex become accessible to binding only when recombinant E2 protein is used in isolation. To present a more native coat protein conformation, we explored availability of the recombinant E1/E2 ectodomain complex. However, the yield and purity were insufficient for the phage selection pipeline (data not shown). These challenges prompted exploration of alternative antigen formats.

**Figure 1.**
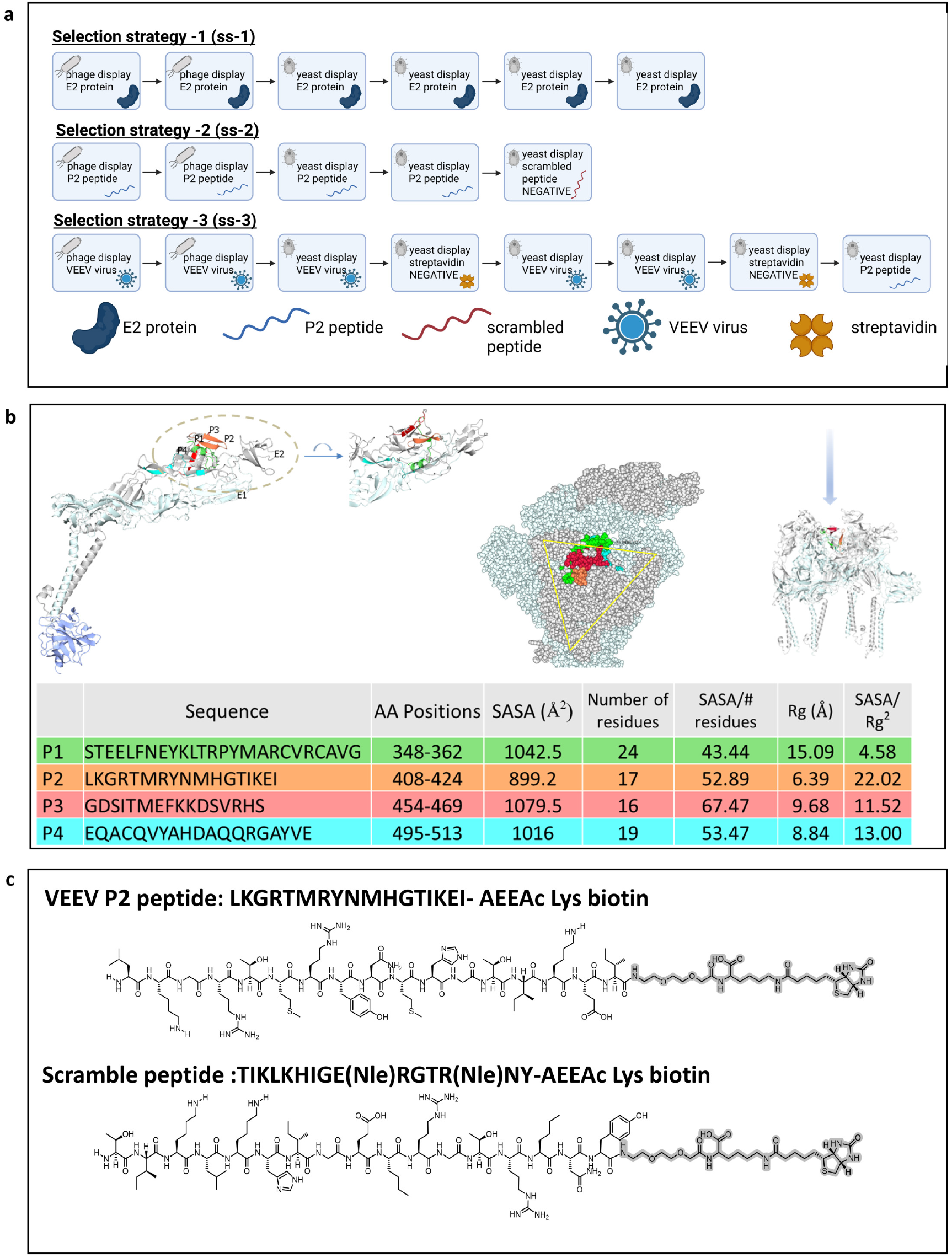
Antigen formats used to select human V_H_Hs to VEEV. Panel A provides a description of selection strategies with recombinant ectodomain (E2), linear peptide, and whole viral particles used in this study, created in BioRender, Velappan, N. (2026) https://BioRender.com/euhrdzl. Panel B provides details of the peptide antigen using computational modeling. Here the E2 fragments mapped onto ribbons representation of E2 (gray) and E1 (light blue) complex. A view of the structure in the dashed circle with 90° rotation is also shown. The fragment IDs and colors correspond to the respective fragments in structure shown above. The right diagram in panel B shows the locations of fragments mapped onto the VEEV virus-like particle structure (PDB:7FFF) looking down the 3-fold axis towards transmembrane helixes. Panel C shows the chemical structure of the P2 peptide, and the corresponding scramble peptide used as competitive selector.

Previously, we demonstrated that antibodies recognizing peptides corresponding to the ectodomain of the influenza A M2 protein can bind the full-length transmembrane protein on cell and viral surfaces.^25^ This finding supported the feasibility of using linear peptides as antigens. Four immunogenic fragments of E2 (P1, P2, P3, and P4), were assessed for nanobody-binding propensity. Peptide locations were mapped to the VEEV structure (PDB IDs: 7FFE/7FFO) as shown in **Fig. 1b**. These peptides are all well-exposed surface fragments since their per-residue solvent accessible surface area (SASA) are all comparable, taking into account the side-chain variability. To semi-quantitatively evaluate surface clustering property (suitable for V_H_H anchoring) of these fragments, we implemented a dimensionless score: SASA / (radius of gyration, Rg^2^). Higher scores represent more congregated patches while lower scores may indicate more elongated or fragmented surfaces. In this test, P2 ranked highest (22.02), and visual inspection confirmed this analysis (**Fig. 1b**). Therefore, P2 was prioritized for further V_H_H antibody selection. Peptide synthesis also offers additional controls, such as unlabeled and scrambled peptides, to enhance specificity during selection pipeline. The peptides were synthesized in good to excellent yields by microwave assisted solid phase syntheses on a CEM Liberty Prime™ using conditions outlined in the materials and methods section. The peptides were initially tested against control single chain variable fraction (scFv) antibody displayed on yeast to examine non-specific interactions against previously characterized antibodies. The norleucine (Nle) version of the P2 peptide was found to bind non-specifically, while the P2 peptide with original sequence and scramble peptide with Nle substitution showed no non-specific interaction during this initial interrogation on antigen quality (**Fig S1b**).

While various forms of the VEEV virus present different opportunities in for antibody selection, there are also potential challenges associated with each form of the target antigen. Recombinant proteins and peptides may differ in folding compared to E1/E2 on the viral surface. For peptides, the termini present in the peptide but not the full version of the protein may participate in the epitope of antibodies when used for selection. As a result, viral peptides may direct antibody binding to a highly specific region, or they have the potential to result in lower or no binding if this C- or N-terminal interaction is needed to mediate binding. On the other hand, the inactivated virus offers the most biologically relevant form of viral proteins, but in this antigen other proteins beyond E2 or the protein of interest are also present, and antibodies may be selected to sites other than the epitope of highest interest. As a result, it is often necessary to either perform alternating selection against different antigens or to check binding against different versions of the target to ensure binding to both the desired site and within the larger viral context. Consequently, we selected sucrose-gradient–purified, UV-inactivated virus as the final antigen format. In particular, the vaccine TC-83 strain of VEEV was used as opposed to the virulent Trinidad Donkey parental strain due to its reduced biosafety constraints and the conserved nature of the glycoproteins.^26^

### Selection of recombinant V_H_Hs against VEEV from a semi-synthetic antibody library

To generate V_H_Hs reactive to the E2 protein of VEEV, we employed a semi-synthetic V_H_H library.^11,17^ Library panning was performed using three distinct antigen formats (**Fig. 1a**). The phage library underwent two rounds of panning, and the phage populations obtained after the second round were evaluated by polyclonal phage ELISA, in which the phage outputs were tested against their corresponding targets (**Fig. 2a-c**). A significant enrichment of specific polyclonal antibody phage occurred after the second round of selection with all three antigen formats. Comparatively, the population selected against the recombinant E2 protein delivered a lower signal in the polyclonal phage ELISA (**Fig 2a**).

**Figure 2.**
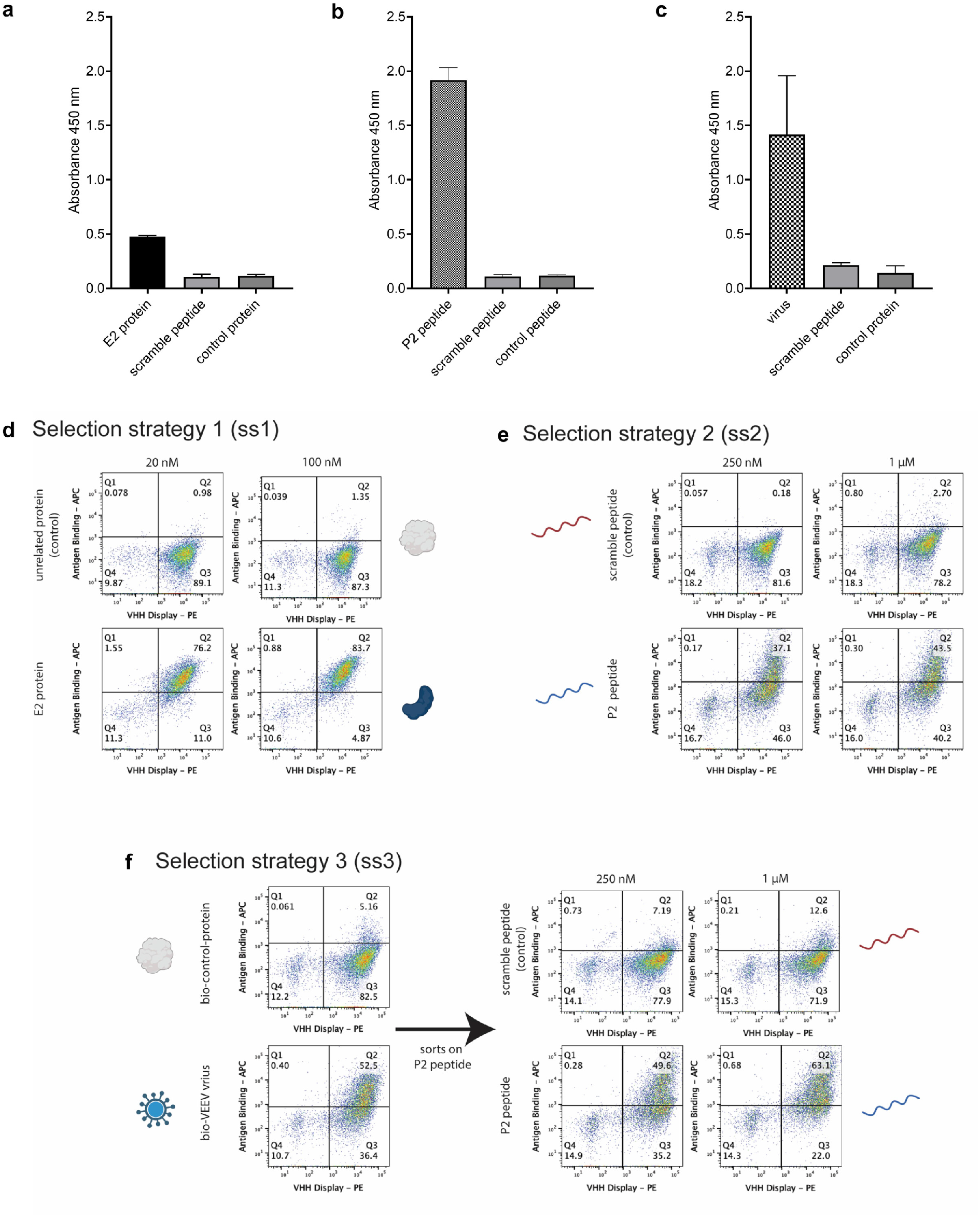
Phage panning and yeast sorting against various VEEV antigens. Binding signal of the polyclonal phage populations obtained after two rounds of phage display selections against a) the biotinylated E2 recombinant protein, b) the VEEV P2 peptide, and c) the biotinylated inactivated viral particles. Analysis of binding signals by flow cytometry of the final antibody population obtained after yeast display against d) the biotinylated E2 recombinant protein, e) the VEEV P2 peptide, and f) the biotinylated inactivated viral particles.

The output selected solely using the full-length E2 protein exhibited a specific yet relatively weak signal against the intended target. In contrast, the peptide-based campaign yielded a robust and highly specific signal for the target peptide, with no detectable reactivity toward the control peptide composed of the same amino acids in a randomized sequence (scramble peptide). Furthermore, inactivated viral particles also demonstrated a strong and specific signal, confirming the recognition of the native antigenic context. The outputs from the second round were subcloned into a yeast display vector and transformed into yeast, following a well-established selection pipeline.^8,19^ During the yeast-based enrichment for binders, we implemented a strategy involving one or two sorts on the same target used during phage selection (either protein, peptide or virus). For the selection on the viral particles, the phage output on yeast contained streptavidin binders (4% of the population) hence a negative sort (collection of population that does not bind to streptavidin) was collected (**Fig. 1A ss-3**). Subsequently, the population was enriched for virus-binders with two rounds of sorting on biotinylated virus, followed by one more round of negative sort on streptavidin binding population. In addition, this population underwent two additional sorts on the peptide (P2) target at 250 nM and 1 µM (**Fig. 2f**) prior to next generation sequencing (NGS) to identify most abundant clones. This approach was designed to ensure that the antibodies selected on the virus would also recognize the epitope of interest in the E2 protein. The final yeast populations (**Fig. 2d-f**) were subjected to sequencing via NGS using Oxford Nanopore Technologies (ONT) to identify the enriched V_H_H clones. The nine most abundant clones identified using the protein as target, the three most abundant clones obtained using the peptide, and the seven most abundant clones obtained when the virus/peptide strategy were selected for further analysis. A total of 19 unique antibodies were gene synthetized, expressed and purified as V_H_H-Fc (human IgG1 Fc) fusion proteins.

All the expressed V_H_H-Fc fusions were assessed in ELISA for their ability to bind the peptide, the E2 protein (**Fig. 3a-c**). ELISA analysis using soluble V_H_H-Fc fusions revealed that all the three clones selected using the VEEV P2 peptide bound to the same peptide, the recombinant E2 protein. In contrast, all the clones selected using solely the recombinant E2 protein were unable to bind the peptide, and only one clone showed a specific binding signal to the E2 protein in ELISA (**Fig. S1a**) but exhibited no binding signal to the viral particles. All the clones obtained from the selection against inactivated biotinylated virus showed binding in ELISA against the peptide, but only one showed binding when tested with the recombinant protein. The V_H_H sequences showed diversity in amino acid sequence, particularly in complementarity-determining region 3. The V_H_Hs were selected from a library based on four clinical scaffolds.^17^ We focused particularly on further characterization of four V_H_Hs (**Fig. 3d**). Of these four, the V_H_Hs selected using only peptide antigen (selection strategy 2), all had the framework region corresponding to clinical antibody sonelokimab. The V_H_H selected using both inactivated virus and peptide had the framework corresponding to clinical antibody caplacizumab.^17^ While these V_H_Hs showed greatest diversity in their CDR3 sequences, they also differed from each other substantially in CDR1. However, all V_H_Hs had similar CDR2 sequences.

**Figure 3.**
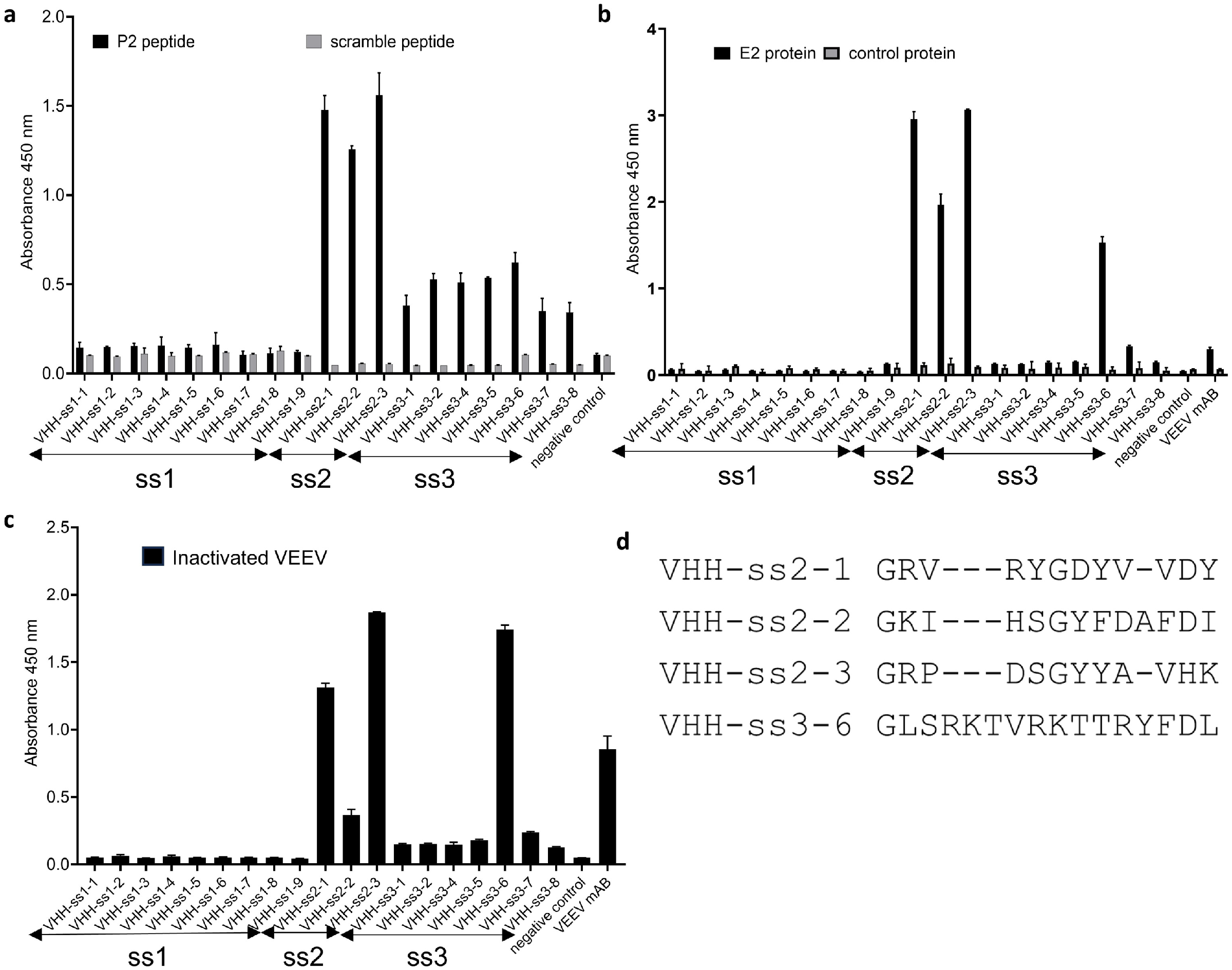
ELISA data from the purified V_H_H Fc-fusions and sequence alignment. Nineteen unique V_H_H Fc-fusions were expressed, purified, and tested against the a) VEEV P2 peptide, b) the E2 recombinant protein, and c) the inactivated viral particles. d) CDR3 alignment for V_H_Hs that recognize the VEEV viral particles.

### Virus binding validations (ELISA, neutralization, and viral sponge properties)

Once a suite of V_H_Hs had been selected, we determined whether they bind to viral particles, indicating recognition of E2 in its native conformation. We coated an ELISA plate with 1×10^7^ PFU/mL VEEV TC-83 and then incubated with 0.25 µg V_H_H-Fc fusion, the commercially available VEEV-57 mAb, or an IgG isotype control.^10^ We identified four of the V_H_H-Fc fusions that bound substantially greater than the IgG control: one antibody obtained from the selection on inactivated viral particles recognizing the peptide in ELISA (VHH-ss3-6) and the three V_H_Hs identified at the end of the selection on the peptide (VHH-ss2-1, VHH-ss2-2 and VHH-ss2-3) (**Fig. 3c)**.

Next, we performed plaque reduction neutralization assays to determine if any of the V_H_Hs could neutralize viral particles in liquid. We incubated 20 µg of V_H_H-Fc fusion or VEEV-57 antibody with 2.5×10^2^ PFU/mL VEEV TC-83 for 1 h, then performed a plaque assay. Mock treated virus (inoculum) served as a negative control. As previously published, all concentrations of the mAb VEEV-57 antibody reduced plaque number and size^10^, however, none of the V_H_H-Fc fusions were found to be neutralizing in solution (**Fig. S2**).

A noteworthy property of V_H_Hs is that they may be readily tethered to other proteins or antibodies, allowing multiple antibodies to be grouped in proximity creating a “sponge” which can be used to bind virions and sequester them in a solution.^27^ Because four of the V_H_Hs were found to bind to VEEV virions (**Fig. 3c**), we wanted to determine if they could act as viral sponges and immunoprecipitate virions out of solution. To test this, we immobilized 20 µg of VHH-ss3-6, VHH-ss2-1, VHH-ss2-2 and VHH-ss2-3, VEEV-57, or IgG isotype control on protein G Dynabeads and then incubated them with 2.5×10^2^ PFU/mL VEEV TC-83 for 1 h. After removing the beads, the supernatant was titered by plaque assay. All the V_H_Hs and the VEEV-57 antibody reduced infectious particles in the supernatant to varying degrees **(Fig. 4)**. VEEV-57 and VHH-ss2-3 demonstrated the greatest efficacy, showing 100% and 98% plaque reduction, respectively, compared to IgG. VHH-ss3-6 reduced plaque formation by 80%, while VHH-ss2-1 showed a 61% reduction, and VHH-ss2-2 showed a 46% reduction.

**Figure 4.**
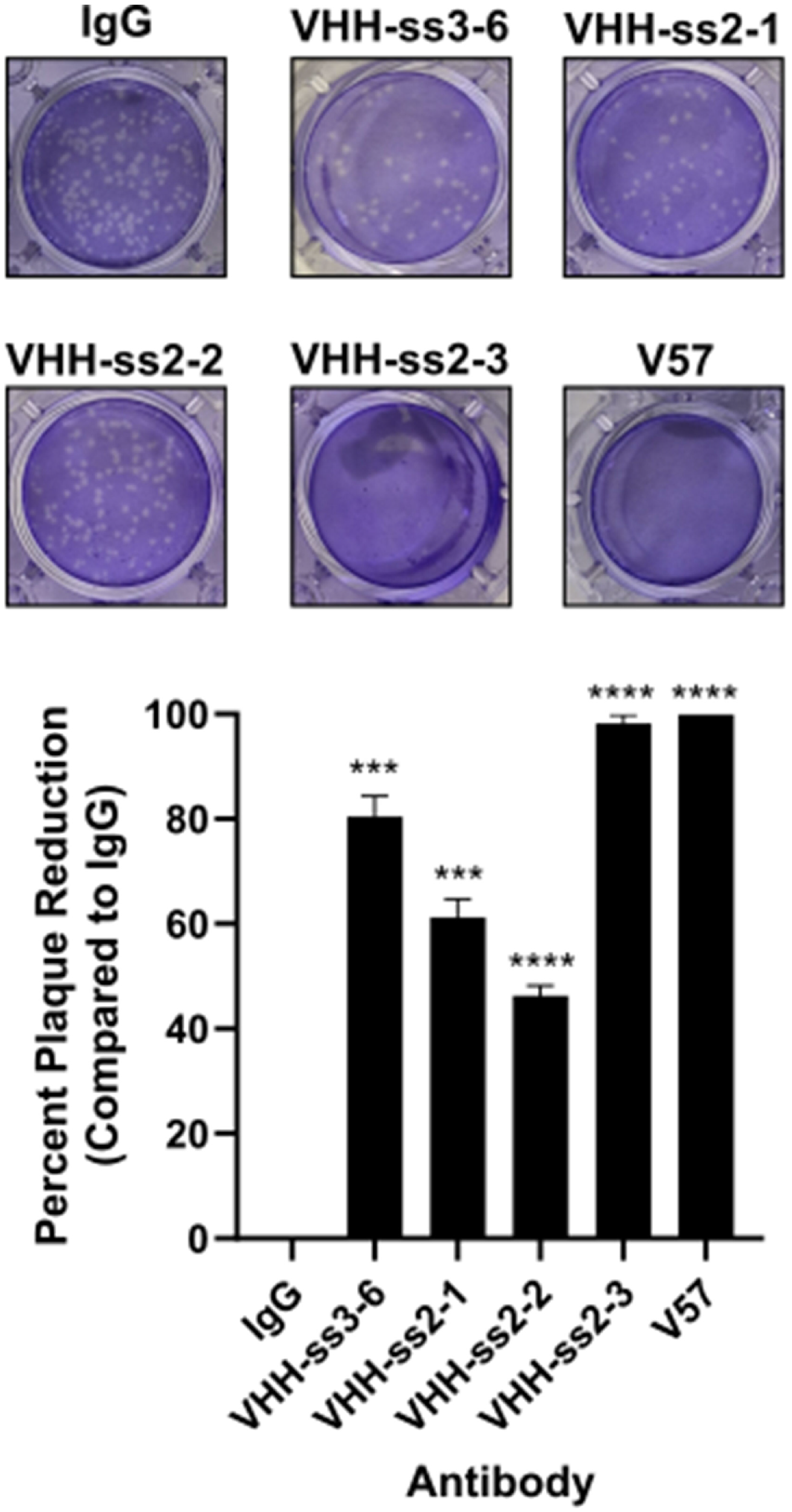
V_H_Hs can act as viral sponges. 20 µg of the indicated V_H_Hs, mAb VEEV-57, or IgG isotype control were incubated with 2.5×10^2^ PFU/mL VEEV virions for 1 h. Antibody-bound viruses were precipitated using Protein G Dynabeads, and viral plaque assays were performed on clarified supernatants. Plaque reduction was calculated as the percentage of plaques compared to IgG (isotype control). Quantitative data shown represents the mean of three technical replicates (+/- SD).Statistical significance was determined by Student’s *t* test using plaque number. ***=p<0.001, ****=p<0.0001

## Discussion

Mosquito-borne diseases remain a significant global health threat, causing hundreds of millions of infections each year^28^. The rising incidence of drug resistance, limited effectiveness of current interventions, and absence of vaccines for several key arboviruses highlight an urgent need for novel, broad-spectrum therapeutics that target both viral and vector-specific mechanisms. In particular, VEEV represents a significant threat to public health due to its propensity for epizootic spillover and potential use as a biothreat. This study reports the screening and isolation of anti-VEEV V_H_Hs from a semisynthetic humanized naïve V_H_H library using phage and yeast display technologies. A key determinant of success in any selection campaign is the choice of target antigen. In this study, we evaluated multiple antigen formats and integrated phage and yeast display to identify V_H_Hs targeting the E2 glycoprotein of VEEV. Our findings demonstrate that antigen presentation significantly shapes the outcome of V_H_H discovery campaigns and that peptide- and virion-based approaches outperform recombinant E2 protein in yielding virus-reactive clones. The purity and availability of well-designed control antigens have proven to be crucial quality requirements for the antigen formats used in phage and yeast display selection processes. Furthermore, our results demonstrate the potential use for non-neutralizing antibodies to be used as viral “sponges”, which could have therapeutic impacts.

Recombinant E2 ectodomain was initially selected as the antigen based on its immunological relevance; however, its requirement for solubilization in urea and subsequent refolding into PBS likely indicate compromised structural integrity. The limited enrichment of phage binders and the weak ELISA reactivity of the resulting V_H_H-Fc fusions support the conclusion that the recombinant antigen did not fully recapitulate the native conformation of E2. The intertwined architecture of the E1/E2 complex, as observed in published alphavirus structures, further suggests that isolated E2 may not adopt the conformations present on the virion surface. Although expression of the E1/E2 heterodimeric ectodomain was attempted, insufficient yield and purity restricted its use in selection campaigns (data not shown).

In contrast, structural mapping and surface accessibility analyses identified a short linear region within E2 (P2 peptide) that is solvent exposed and part of the known antigenic epitope for VEEV.^24^ Phage selections using this peptide produced strong enrichment, high specificity relative to scrambled control, and clones that recognized the peptide, recombinant E2, and inactivated virions. These results align with earlier observations^25^ that antibodies raised against short extracellular peptides of viral membrane proteins can retain reactivity to the intact virus. The ability of P2-selected V_H_Hs to bind multiple antigen formats indicates that this peptide effectively mimics a naturally accessible linear epitope on the virion surface.

Selections performed using inactivated viral particles yielded a second set of V_H_Hs with strong virion reactivity compared to recombinant protein selections. Most virus-selected clones also recognized the P2 peptide, indicating convergence on the same immunodominant region identified structurally and through peptide-based selection. However, only one virus-derived clone bound the recombinant protein, again underscoring discrepancies between recombinant E2 and its native viral context.

Neutralizing antibodies typically block viral entry to host cells by physically blocking binding to host receptor, often facilitated by the binding angle and the Fc mediated steric hindrance.^29^ This blockage reduces the infection rate and lowers the viral titer. Neutralizing antibodies typically bind to highly variable epitopes such as the receptor binding domain of the SARS-CoV-2 spike protein.^30^ These epitopes undergo mutations as the virus tries to evade host immune response, reducing the efficacy of these antibodies as new variants arise. A strategy used in this scenario is developing broadly neutralizing antibodies that bind to conserved regions, however identification of such epitopes for emerging pathogens may take extensive time and research. Non-neutralizing (nNAbs) antibodies have previously been isolated from immunization with multiple viral species, and their effect in lowering viral load has been demonstrated.^31^ Known mechanisms of action for nNAbs include antibody dependent cytotoxicity (ADCC), complement-mediated cytotoxicity, steric inhibition of proteins important in viral replication, and antibody-dependent cell-mediated phagocytosis (ADCP), which are mediated by the Fc region of antibodies.

Although four V_H_Hs demonstrated binding to immobilized virions, none neutralized the virus in plaque reduction assays. These findings suggest that the targeted epitope, while accessible and immunodominant, is not essential for entry-related functions. The robust neutralization observed with the positive-control mAb VEEV-57 confirms assay sensitivity and highlights the functional distinction between high-affinity binding and neutralizing capacity. It should be noted that the peptides used for development of these V_H_Hs are in not in proximity of mutation sites (**Fig. 4**), including the major virulence determinant at aa120, that differ between TC-83 and the virulent Trinidad donkey strain.^32,33^ Validation of binding to virulent VEEV strains under BSL-3 conditions in future studies would further clarify the utility of the V_H_Hs described herein.

Despite the absence of neutralization, all virus-binding V_H_Hs effectively removed infectious virions from solution when immobilized on protein G Dynabeads. This “viral sponge” activity demonstrates that these V_H_Hs can sequester virus with high efficiency, with one clone approaching the performance of mouse monoclonal VEEV-57 in reducing infectious titers. These data indicate that V_H_Hs targeting non-neutralizing epitopes can still perform valuable functions in virion capture, concentration, and removal applications. Further work is needed to determine the therapeutic potential of V_H_H viral “sponges”.

An advantage of non-neutralizing antibodies and V_H_Hs is that they can be functionalized and/or modified to increase their utility while taking advantage of the lack of viral evolution directed towards their epitopes. Popular mechanisms explored include modifications or switching the Fc domain to increase the Fc effector function or creation of multivalent antibodies to increase binding avidity.^14^ In addition to Fc domain, V_H_Hs can also be functionalized using multimerizing units (e.g., multimeric proteins, self-assembling nanomaterials) to facilitate integration of favorable characteristics to further improve potential therapeutics against viral pathogens, especially emerging viruses with pandemic potential. A key advantage of V_H_Hs is the ability to tether them with compatible materials to increase their functionality. The viral sponge property of these nanobodies could be improved by combining them as bispecific or multispecific binders. Self-assembling biopolymers is an example of suitable material for these purposes.

Anti-VEEV V_H_Hs isolated from immunized of llamas followed by phage display screening have been characterized previously.^5,6^ These single domain nanobodies when presented in bivalent format provide viral neutralization *in vitro* and *in vivo* and structural analysis of binding epitopes reveal that these V_H_Hs bind in close proximity to the LDLRAD3 receptor binding site.^34^ In contrast, the V_H_Hs described herein have framework region and CDRs derived from human antibodies. In addition, the P2 peptide used as the target antigen in our selections present an orthogonal viral epitope (**Fig. S4**) to the published animal derived moieties. Together, these differences suggest the potential for combining these V_H_Hs with existing molecules to enhance therapeutic efficacy in future studies.

Overall, this work demonstrates that antigen format is a critical determinant of success in V_H_H discovery against structurally complex viral glycoproteins. Peptide-based and virion-based campaigns converged on a shared, surface-exposed epitope and yielded V_H_Hs with robust virion-binding and viral capture properties. Future studies will benefit from exploring additional epitopes beyond P2, applying affinity maturation, and evaluating multivalent or Fc-extended constructs that may enhance neutralizing potential. This antigen-selection framework may be broadly applicable to other enveloped viruses with conformationally sensitive surface proteins. Furthermore, the human V_H_Hs described in this study represent a promising medical countermeasure (MCM) platform with potential for rapid deployment and stockpiling. Specifically, the anti-VEEV V_H_Hs developed here with the ability to act as viral sponges could serve as a valuable foundation for MCM preparedness strategies aimed at protecting deployed personnel and vulnerable populations against emerging mosquito-borne viral threats.

## Materials and Methods

### Viruses and mammalian cells

VEEV TC-83 was obtained through BEI Resources, NIAID, NIH: Venezuelan Equine Encephalitis Virus, TC-83, NR-63. Vero cells (ATCC, CRL-81) were cultured in Dulbecco’s modified Eagle’s medium (DMEM, VWR 45000-304) supplemented with 1% (v/v) nonessential amino acids 100x solution (Gibco/Fisher Scientific 11140050), 1% (v/v) Penicillin Streptomycin 100x Solution (Corning, 30-002-Cl), 2.5% HEPES (Gibco/Fisher Scientific 15-630-106), and 10% heat-inactivated FBS. Cell lines were cultured at 37°C and 5% CO_2_ in a humidified incubator. Regular passaging using standard subculturing techniques was used to maintain low-passage-number stocks.

### Bacteria and yeast strains

OmniMAX™ 2 T1R *E. coli* (ThermoFisher, C854003): F’ [proAB+lacIqlacZΔM15 Tn10(TetR) Δ(ccdAB)] mcrA Δ(mrr-hsdRMS-mcrBC) φ80lacZΔM15 Δ(lacZYA-argF)U169 endA1 recA1 supE44 thi-1 gyrA96 (NalR) relA1 tonA panD was used for phage display. *S. cerevisiae* EBY100 (ATCC, MYA-4941): (GAL1-AGA1::URA3 ura3-52 trp1 leu21 his3200 pep4::HIS2 prb11.6R can1 GAL) was used for yeast display

### Antigen preparation

#### Protein antigen

Purified VEEV E2 glycoprotein (Creative Diagnostics, DAGA-268) was purchased and resuspended in 1xPBS with 8M Urea per manufacturer’s instruction and then dialyzed into 1xPBS prior to biotinylation using EZ-Link™ Sulfo NHS-LC-LC-Biotin (ThermoFisher, 21343) following manufacturer’s instructions.

#### Peptide antigens

Six different peptides were synthesized (**Fig. S3**). The biotin labeled peptides were used as antigens in phage and yeast selections and the unlabeled peptides were synthesized for use as competition during the selection process.

##### Computational Modeling

SASA of selected E2 fragments were calculated using the BIO.PDB.SASA module in the BioPython package. Rg values were analyzed with the HullRad software^35^ using the coordinates of the corresponding fragments extracted from the VEEV structure (PDBID:7FFO). The Ribbon diagram in **Fig 1b** was generated with the CCP4mg^36^ package.

##### Automated Solid Phase Peptide Synthesis (SPPS)

A CEM Liberty Prime™ microwave peptide synthesizer was used for solid phase synthesis at high temperature (90 °C). All syntheses were performed at a 0.1 mmol scale using the recommended standard instrument chemistry, with modified cycles in which drain times were increased from 5 to 10 s to accommodate for any resin volume increases over successive synthesis cycles. Coupling of Fmoc-protected amino acids (5 eq, 0.5 M in DMF) was achieved by treatment with DIC (10 eq, 2 M in DMF) and Oxyma Pure (5 eq, 0.25 M in DMF). For the peptides described in this study, double-coupling instrument cycles were employed, resulting in average coupling yields of >98% per cycle. The synthesis deploys direct addition of 25% pyrrolidine in DMF to the heated resin for Fmoc deprotection at 105 °C for 60 s, a 120 s coupling at 90 °C, and three wash steps of 20 s each, resulting in a total single coupling cycle time of 4 min. To prevent hydrolysis of acid-labile side-chain protecting groups during the syntheses, 0.1 M DIPEA was added to the Oxyma solution.

##### C-terminal Biotinylation

Peptides bearing a *C*-terminal biotin were synthesized on Fmoc-Lys(Biotin)-Wang resin using standard Fmoc SPPS. The preloaded biotinylated lysine served as the *C*-terminal residue. After resin swelling, the *N*-terminal Fmoc group was removed with 25% pyrrolidine in DMF, and the Fmoc-AEEAC-OH linker was installed using DIC/Oxyma activation. Then, the remaining peptide sequence was assembled by iterative cycles of Fmoc deprotection and amino acid coupling, also using DIC/Oxyma activation. Following completion of the sequence, peptides were cleaved from resin and globally deprotected under standard TFA cleavage conditions. The biotin moiety is stable in these conditions, yielding the peptides with a biotin label.

##### Peptide Cleavage and Global Deprotection

Deprotection used 25 mL of modified “reagent K” mixture consisting of TIPS (1.25 mL), thioanisole (0.625 mL), DODT (1.25 mL) - a less odorous substitute for EDT (ethylendithiol), water (1.25 mL), and TFA (trifluoroacetic acid). The resin was pretreated with the quencher solution for 5 min, then the TFA was added (to a final volume of 25 mL). The deprotections were carried out in 50 mL conical tubes of high-density polypropylene, under a blanket of Argon to prevent side reactions from air for 3 h at room temperature. The solutions were filtered, and the filtrate was concentrated to 10 mL. The peptide was then precipitated into ice cold ether and collected by centrifugation.

##### Purification and Analysis of Synthetic Peptides

Purifications (to > 98%) were performed on a Waters HPLC preparative work-station with a 2545 pump, at a flow rate of 20 mL/min. C18 reverse-phase columns (Waters BEH (Ethylene Bridged Hybrid) 130 Å, 10 μm, 19×150 mm) with a gradient from water to acetonitrile with 0.1% TFA added. Peaks were collected based on monitoring at 215 nm using a PDA 2998 detector. Combined product fractions were lyophilized, yielding white fluffy solids. Reaction yields were between 70 and 90%, indicating coupling yields of > 99% per amino residue incorporated. Peptides were then analyzed for purity by analytical HPLC on a C18 reverse phase column (Waters BEH 130 Å, 5 μm, 4.6×150 mm) with a linear gradient from 95% to 50% acetonitrile water with 0.1% TFA in a 10 min gradient, and by mass spectrometry on a Thermo LTQ mass spectrometer in ESI+ mode.

#### Inactivated whole virus as antigen

To propagate the VEEV TC-83 stocks, 5×10^6^ Vero E6 cells were infected with VEEV TC-83 at an MOI of 0.01 PFU/cell. Supernatants were collected at 3 days post-infection and clarified by centrifuging at 1000g for 10 minutes. Virus used as antigen for antibody production was inactivated by applying UV radiation at 2500 mJ/cm^2^ (FisherBrand UV crosslinker, 13-245-221). The inactivated viral stock was then pelleted though a 30% w/v sucrose cushion in TNE buffer (10 mM Tris-HCl pH 7.5, 100 mM NaCl, 1 mM EDTA) for 2 h at 30,000 rpm and resuspended in 100 µL PBS. The viral particles were biotinylated similar to the protein antigens as described above.

### V_H_H selections

For phage selections, 10 µL of streptavidin-conjugated magnetic beads (Dynabeads M-280, ThermoFisher, 11205D) were coated with an excess of biotinylated antigen to ensure complete coverage. The selection process was performed using the automated KingFisher magnetic bead system (ThermoFisher Scientific), which included multiple washing steps to remove unbound phage. Bound phage were eluted from the beads by acid treatment and used to infect F′ pilus-carrying bacteria. After propagation of the eluted phage, the selection cycle was repeated. After two rounds of phage selection, the enriched V_H_Hs were subcloned into the yeast display vector as previously described.^19,37,38^ The selected V_H_H genes were amplified using primers that introduced overlapping sequences to the yeast display vector pDNL6. The vector and amplified fragments were co-transformed into yeast cells to enable cloning by homologous recombination.^8,19^ The resulting yeast mini-libraries were further enriched for binders using fluorescence-activated cell sorting following established protocols.^19,37,38^ After induction, 2 × 10^6^ yeast cells were stained with 100 nM biotinylated antigens or 10^6^ biotinylated viral particles. Cells were then labeled with streptavidin–AlexaFluor633 (ThermoFisher, S21375) to detect antigen binding and anti-SV5-phycoerythrin (PE) to assess V_H_H display levels. Yeast clones positive for both antigen binding (AlexaFluor633) and display (PE) were sorted. Sorted cells were grown at 30 °C for two days and induced for the next round of sorting at 20 °C for 16 h.

### ONT Sequencing and VHH-Fc expression

Yeast plasmid DNA was purified from the final saturated yeast cultures using a Qiagen miniprep kit (Qiagen, 27106) with the addition of acid-washed glass beads and vortexing to promote lysis. Purified DNA was eluted in 50 µL Buffer EB. To amplify the cloned V_H_H for NGS, 100 µL PCR reactions were prepared each with 25 µL of yeast miniprep as template, 0.2 µM forward and reverse primers, and 1 unit NEB Q5 Hot-Start Polymerase. The same forward and reverse primers were used as for Sanger sequencing. The reactions were cycled at 98 °C for 30 s; 20 cycles of 98 °C for 5 s, 65 °C for 5 s, and 72 °C for 25 s for V_H_H; 72 °C for 2 min. The amplified V_H_H (around 640 bp) were purified from a 1% agarose gel and submitted to Plasmidsaurus^®^ for sequencing using the Standard Premium PCR Service. The antibody sequences were submitted to Biointron^®^ for gene synthesis, cloning and protein expression as V_H_H humanIgG1 Fc-fusion, and purification.

### V_H_H validation

#### ELISA

Nunc-Immuno™ MicroWell™ 96 well MaxiSorp™ (ThermoFisher, 442404) were coated overnight with neutravidin (ThermoFisher, 31050, 2.5 µg/mL). The following day, after removing the neutravidin solution, 2.5 µg/mL of biotinylated-E2, biotinylated-target peptide, biotinylated-scrambled peptide, and biotinylated-interferon gamma (IFNγ) (negative control), were added to the plate. After binding, wells were washed three times with 1x PBS and blocked with 5% bovine serum albumin (BSA) at room temperature for 1 h. The purified V_H_H-Fc (100 µL) were added to the wells at a concentration of 1 µg/mL in 1x PBS and incubated for 1 h. Wells were washed three times with PBS and 100 µL of anti-human IgG (1:2000 in 1x PBS 0.5% BSA, Jackson ImmunoResearch, 109–035–008) were added per well. After 1 h incubation at room temperature, wells were washed three times with 1x PBS and 50 µL of 3,3’, 5,5’ tetramethylbenzidine (TMB) substrate (Sigma, T8665-1 L) were added per well. To stop the reactions, 25 µL of 1 M sulfuric acid (H_2_SO_4_) were added per well. Absorbance was read at 450 nm using a Tecan Infinite 200 PRO plate reader.

Inactivated VEEV TC-83 was diluted to 1×10^7^ PFU/mL to coat 96-well ELISA plates (ThermoFisher Scientific, 3455) with 100 µL/well and incubated overnight at 4°C. The plates were then blocked for 1 h at room temperature with 5% FBS in PBS. V_H_H-Fc fusions, VEEV-57 mAb (BioXCell, BE0435), or IgG isotype control (MilliporeSigma, M5284) were diluted to 2.5 µg/mL in 5% FBS, added at a volume of 50 µL per well, and incubated for 2 h at room temperature. Plates were washed three times with PBS and then secondary antibody, either goat anti-human (V_H_H; Jackson ImmunoResearch 109-035-088) or goat anti-mouse (VEEV-57 or IgG; Jackson ImmunoResearch, 115-035-003), was diluted 1:5000 in 5% FBS and incubated for 2 hat room temperature. The plates were then washed three times with PBS and 100 µL TMB solution was added per well and incubated 5 min, then the reaction was stopped with 1% HCl. Absorbance was then measured at OD 450nm with a Synergy HTX plate reader.

#### Plaque Reduction Neutralization Titer Assay

V_H_H (15 µg/mL) or VEEV-57 (ranging from 2.5 µg/mL-10 µg/mL) antibodies were incubated with 2.5 x 10^2^ PFU/mL VEEV TC-83 at 37°C for 1 h. The antibody-virus mixture was then added to 80% confluent Vero cells and allowed to adsorb for 1 h at 37°C. A 10-fold serial dilution of the inoculum without antibody was used to calculate inoculum titer and as a positive control. Afterwards, cells were overlaid with a solution of 1.2% cellulose (Sigma Aldrich, 435244-250G) in 1x complete DMEM medium for 4 days post infection (dpi). The monolayers were then fixed with formaldehyde solution (4% formaldehyde-1x PBS) for at least 1 h. The overlay was then removed, and the plaques were visualized via crystal violet staining.

#### VEEV Immunoprecipitation Assay

20 µg of V_H_H, VEEV-57, or IgG (Millipore Sigma,) isotype control antibody were bound to 50 µL of Protein G Dynabeads (ThermoFisher, 10004D) in 500 µL PBS and rotated at room temperature for 2 h. 2.5×10^2^ PFU/mL VEEV TC-83 was then added to the antibody bound beads and the mixture was rotated at 37 °C for 1h. The beads were then immobilized on a magnet, and the supernatant was added to 80% confluent Vero cells and allowed to adsorb for 1 h at 37 °C. Afterwards, cells were overlaid with a solution of 1.2% cellulose in 1x complete DMEM medium for 4 days post-infection. The monolayers were then fixed with formaldehyde solution (4% formaldehyde-1x PBS) for at least 1 h. The overlay was then removed, and the plaques were visualized via crystal violet staining.

## Acknowledgements

This project was funded by a grant from Defense Threat Reduction Agency (award# CB11435 to NV). This manuscript has been reviewed at Los Alamos National Laboratory and assigned report number LA-UR-26-20046. The authors would like to thank Dr. Tiffany Nguyen for her support of this work. ATL was supported by a fellowship from the National Institutes of Health NIGMS T32AI007538. We acknowledge that the VEEV virus was obtained through BEI Resources, NIAID, NIH: Venezuelan Equine Encephalitis Virus, TC-83, NR-63.

## Author Contributions

Conceptualization – NV, JMZ, JS; Experimental methodology – ATL, FF, JS, JMZ, ALM, NV; Computation modeling - LH; Writing - original draft preparation, ATL, FF, JMZ, JS, LH, NV; Critical review - all authors, Supervision – AMK, NV.; Project administration, NV.; Funding acquisition, NV.

---

**Supplementary Figure 1. Binding assay with V_H_Hs selected on recombinant E2 protein. a)** In this ELISA-based assay, the V_H_H Fc-fusion proteins were coated on maxisorp plates at 10µg/mL concentration in 1x PBS. The biotinylated E2 protein bound to the antibodies was detected using streptavidin-PE conjugate. This alternate orientation ELISA, when the protein was allowed to bind in the same format as the yeast display sorting conditions, showed small level of specific recognition for the recombinant E2 protein compared to the first tested orientation (Fig. 3). b) Peptide non-specific analysis on control yeast population is shown, indicating lack of hydrophobic interaction for the native P2 peptide and high level of “stickiness” for the P2 peptide where methionine was substituted with norleucine (Fig. S3).

**Supplementary Figure 2. VEEV V_H_H plaque reduction neutralization assay.** VEEV TC-83 was incubated with either 15 µg/mL of each V_H_H or a range of 1.25 µg/mL – 10 µg/mL of mAb VEEV-57 at 2.5×10^2^ PFU/mL for 1 h. Each virus-antibody mixture was then used to infect Vero E6 cells for a plaque assay. Three technical replicates were performed, and images show representative results.

**Supplementary Figure 3. Sequence of the P2 peptide and derivatives.** Three sets of peptides were synthesized. The first set includes the native P2 peptide that is present in the E2 protein of VEEV along with its biotin-labeled version used in the selections. The second set contains the same peptide sequences in which methionine was substituted with Norleucine to prevent oxidation of the sulfur atom (i.e., formation of methionine sulfoxide or methionine sulfone). The last set shows sequences for scramble peptides that were used as control in yeast sorting.

**Supplementary Figure 4. P2 peptide epitope information.** Computational model of the E1 (light blue) and E2 (grey) complex indicating the P1, P2, and P3 peptides (green, orange, red respectively) are shown along with known amino acid changes (yellow circles) that differ between the TC-83 and TRD strains of VEEV. In addition, the figure also shows the contact residues (magenta circles) for one of the single domain antibodies (V3A8f) previously described. Panel B shows the same image at 90° rotation angle.

## References

1. Sissoko, D. et al. Post-epidemic Chikungunya disease on Reunion Island: course of rheumatic manifestations and associated factors over a 15-month period. PLoS Negl Trop Dis 3, e389 (2009). 10.1371/journal.pntd.0000389

2. Schwartz, O. & Albert, M. L. Biology and pathogenesis of chikungunya virus. Nat Rev Microbiol 8, 491–500 (2010). 10.1038/nrmicro2368

3. Calisher, C. H. Medically important arboviruses of the United States and Canada. Clin Microbiol Rev 7, 89–116 (1994). 10.1128/cmr.7.1.89

4. de la Monte, S., Castro, F., Bonilla, N. J., Gaskin de Urdaneta, A. & Hutchins, G. M. The systemic pathology of Venezuelan equine encephalitis virus infection in humans. Am J Trop Med Hyg 34, 194–202 (1985). 10.4269/ajtmh.1985.34.194

5. Gardner, C. L. et al. Demonstration of in vivo efficacy, cryo-EM-epitope identification, and breadth of two anti-alphavirus bispecific single domain antibodies. J Virol, e0187525 (2025). 10.1128/jvi.01875-25

6. Liu, J. L. et al. Bivalent single domain antibody constructs for effective neutralization of Venezuelan equine encephalitis. Sci Rep 12, 700 (2022). 10.1038/s41598-021-04434-x

7. Focosi, D. et al. Monoclonal antibody therapies against SARS-CoV-2. The Lancet Infectious Diseases 22, e311–e326 (2022). 10.1016/S1473-3099(22)00311-5

8. Ferrara, F. et al. A pandemic-enabled comparison of discovery platforms demonstrates a naïve antibody library can match the best immune-sourced antibodies. Nature Communications 13, 462 (2022). 10.1038/s41467-021-27799-z

9. Velappan, N. et al. Healthy humans can be a source of antibodies countering COVID-19. Bioengineered 13, 12598–12624 (2022). 10.1080/21655979.2022.2076390

10. Kafai, N. M. et al. Neutralizing antibodies protect mice against Venezuelan equine encephalitis virus aerosol challenge. J Exp Med 219 (2022). 10.1084/jem.20212532

11. Alexander, E. & Leong, K. W. Discovery of nanobodies: a comprehensive review of their applications and potential over the past five years. Journal of Nanobiotechnology 22, 661 (2024). 10.1186/s12951-024-02900-y

12. Zupancic, J. M. et al. Engineered Multivalent Nanobodies Potently and Broadly Neutralize SARS-CoV-2 Variants. Adv Ther (Weinh) 4, 2100099 (2021). 10.1002/adtp.202100099

13. van der Linden, R. H. J. et al. Comparison of physical chemical properties of llama VHH antibody fragments and mouse monoclonal antibodies. Biochimica et Biophysica Acta (BBA) - Protein Structure and Molecular Enzymology 1431, 37–46 (1999). 10.1016/S0167-4838(99)00030-8

14. Mullin, M. et al. Applications and challenges in designing VHH-based bispecific antibodies: leveraging machine learning solutions. mAbs 16, 2341443 (2024). 10.1080/19420862.2024.2341443

15. McMahon, C. et al. Yeast surface display platform for rapid discovery of conformationally selective nanobodies. Nat Struct Mol Biol 25, 289–296 (2018). 10.1038/s41594-018-0028-6

16. Nakakido, M., Kinoshita, S. & Tsumoto, K. Development of novel humanized VHH synthetic libraries based on physicochemical analyses. Scientific Reports 14, 19533 (2024). 10.1038/s41598-024-70513-4

17. Erasmus, M. F. et al. Developing drug-like single-domain antibodies (VHH) from in vitro libraries. MAbs 17, 2516676 (2025). 10.1080/19420862.2025.2516676

18. Zhang, W. et al. Developability assessment at early-stage discovery to enable development of antibody-derived therapeutics. Antibody Therapeutics 6, 13–29 (2022). 10.1093/abt/tbac029

19. Ferrara, F. et al. Recombinant renewable polyclonal antibodies. mAbs 7, 32–41 (2015). 10.4161/19420862.2015.989047

20. Zupancic, J. M. et al. Directed evolution of potent neutralizing nanobodies against SARS-CoV-2 using CDR-swapping mutagenesis. Cell Chem Biol 28, 1379-1388.e1377 (2021). 10.1016/j.chembiol.2021.05.019

21. Velappan, N. et al. Selection and characterization of FcεRI phospho-ITAM specific antibodies. MAbs 11, 1206–1218 (2019). 10.1080/19420862.2019.1632113

22. Velappan, N. et al. Selection and verification of antibodies against the cytoplasmic domain of M2 of influenza, a transmembrane protein. MAbs 12, 1843754 (2020). 10.1080/19420862.2020.1843754

23. Zhang, R. et al. 4.4 Å cryo-EM structure of an enveloped alphavirus Venezuelan equine encephalitis virus. The EMBO Journal 30, 3854-3863-3863 (2011). 10.1038/emboj.2011.261

24. Schein, C. H. et al. PCP consensus protein/peptide alphavirus antigens stimulate broad spectrum neutralizing antibodies. Peptides 157, 170844 (2022). 10.1016/j.peptides.2022.170844

25. Gabbard, J. et al. A humanized anti-M2 scFv shows protective in vitro activity against influenza. Protein Eng Des Sel 22, 189–198 (2009). 10.1093/protein/gzn070

26. Kinney, R. M., Johnson, B. J., Welch, J. B., Tsuchiya, K. R. & Trent, D. W. The full-length nucleotide sequences of the virulent Trinidad donkey strain of Venezuelan equine encephalitis virus and its attenuated vaccine derivative, strain TC-83. Virology 170, 19–30 (1989). 10.1016/0042-6822(89)90347-4

27. Burton, D. R. Antiviral neutralizing antibodies: from in vitro to in vivo activity. Nature Reviews Immunology 23, 720–734 (2023). 10.1038/s41577-023-00858-w

28. Franklinos, L. H. V., Jones, K. E., Redding, D. W. & Abubakar, I. The effect of global change on mosquito-borne disease. The Lancet Infectious Diseases 19, e302–e312 (2019). 10.1016/S1473-3099(19)30161-6

29. Mader, K. & Dustin, L. B. Beyond bNAbs: Uses, Risks, and Opportunities for Therapeutic Application of Non-Neutralising Antibodies in Viral Infection. Antibodies (Basel) 13 (2024). 10.3390/antib13020028

30. Izadi, A. & Nordenfelt, P. Protective non-neutralizing SARS-CoV-2 monoclonal antibodies. Trends in Immunology 45, 609–624 (2024). 10.1016/j.it.2024.06.003

31. Excler, J. L., Ake, J., Robb, M. L., Kim, J. H. & Plotkin, S. A. Nonneutralizing functional antibodies: a new “old” paradigm for HIV vaccines. Clin Vaccine Immunol 21, 1023–1036 (2014). 10.1128/cvi.00230-14

32. Kinney, R. M. et al. Attenuation of Venezuelan equine encephalitis virus strain TC-83 is encoded by the 5’-noncoding region and the E2 envelope glycoprotein. J Virol 67, 1269–1277 (1993). 10.1128/jvi.67.3.1269-1277.1993

33. Johnson, B. J., Kinney, R. M., Kost, C. L. & Trent, D. W. Molecular determinants of alphavirus neurovirulence: nucleotide and deduced protein sequence changes during attenuation of Venezuelan equine encephalitis virus. J Gen Virol 67 ( Pt 9), 1951–1960 (1986). 10.1099/0022-1317-67-9-1951

34. Basore, K. et al. Structure of Venezuelan equine encephalitis virus in complex with the LDLRAD3 receptor. Nature 598, 672–676 (2021). 10.1038/s41586-021-03963-9

35. Fleming, P. J. & Fleming, K. G. HullRad: Fast Calculations of Folded and Disordered Protein and Nucleic Acid Hydrodynamic Properties. Biophys J 114, 856–869 (2018). 10.1016/j.bpj.2018.01.002

36. McNicholas, S., Potterton, E., Wilson, K. S. & Noble, M. E. Presenting your structures: the CCP4mg molecular-graphics software. Acta Crystallogr D Biol Crystallogr 67, 386–394 (2011). 10.1107/s0907444911007281

37. Ferrara, F. et al. Using phage and yeast display to select hundreds of monoclonal antibodies: application to antigen 85, a tuberculosis biomarker. PLoS One 7, e49535 (2012). 10.1371/journal.pone.0049535

38. Azevedo Reis Teixeira, A. et al. Drug-like antibodies with high affinity, diversity and developability directly from next-generation antibody libraries. MAbs 13, 1980942 (2021). 10.1080/19420862.2021.1980942

